# Dissecting Cold Tolerance in *Drosophila ananassae*: A Multi-Phenotypic and Bulk Segregant Analysis

**DOI:** 10.1101/2025.04.23.650207

**Authors:** Vera Miyase Yılmaz, Vincent Ohlhauser, Fikri Tuğberk Kara, Sonja Grath

## Abstract

Temperature strongly shapes the geographic distribution of insects, making cold tolerance an important component of adaptation to different thermal environments. Previous work in *Drosophila ananassae* identified variation in chill coma recovery time both within an ancestral Bangkok population and between this population and a derived Kathmandu population. However, cold tolerance was characterized only by chill coma recovery time, leaving open whether variation in this phenotype reflects cold tolerance more broadly.

Here, we further characterized cold tolerance in iso-female lines from the Bangkok and Kathmandu populations and in recombinant inbred lines generated from the ancestral population, using lethal time and cold shock mortality in addition to chill coma recovery time. We showed that cold tolerance phenotypes differ between sexes, the three phenotypes showed partially overlapping but distinct patterns, and some recombinant inbred lines had extreme phenotypes with higher tolerance than the tolerant founder or lower tolerance than the sensitive founder. We performed bulk segregant analysis using the recombinant inbred lines that exhibited extreme chill coma recovery time to identify genomic regions associated with variation in this phenotype. We identified 22 candidate regions associated with variation in chill coma recovery time and showed that the genes in the putative regions were enriched in muscle development, metabolic processes, and cytoskeletal protein binding. Our results highlight the importance of considering multiple cold tolerance phenotypes and provide insights into the genetic architecture of cold tolerance variation in natural populations of *Drosophila*.

**ARTICLE SUMMARY:** Cold tolerance is often treated as a single trait, but different measures of cold tolerance may reflect different aspects of an organism’s response to low temperatures. Using natural populations and recombinant inbred lines of the vinegar fly *Drosophila ananassae*, we show that commonly used cold tolerance phenotypes capture partially overlapping but distinct aspects of cold tolerance. Recombinant inbred lines show substantial phenotypic variation, and bulk segregant analysis identifies multiple candidate genomic regions associated with variation in chill coma recovery time. Our results highlight the complexity of cold tolerance and its underlying genetic architecture.

## INTRODUCTION

Insects occupy a wide range of ecosystems across the globe and must cope with diverse environmental conditions. Among the many abiotic factors shaping their distribution, temperature is one of the most important and variable components of the environment. Adaptation to thermal conditions is therefore essential for survival and can also facilitate range expansion. Insects employ various physiological and behavioral strategies to survive temperatures outside their optimal range, allowing them to avoid freezing or tolerate chilling (Baust & Lee, 1987; Ring, 1982; Teets et al., 2008, 2012; Teets & Denlinger, 2013; Wang et al., 2017). However, prolonged exposure to low temperatures can cause injury and mortality in chill-susceptible insects (Sinclair, 1999). Disentangling the genetic basis of variation in cold tolerance is therefore critical for understanding thermal adaptation.

In chill-susceptible insects, exposure to low temperatures disrupts ion and water homeostasis and leads to chilling injuries (Sinclair, 1999; Strachan et al., 2011; MacMillan et al., 2015). When temperatures approach the critical thermal minimum (CTmin), insects lose neuromuscular coordination and enter a reversible chill coma. Chill coma recovery time (CCRT), measured after flies are returned to warmer temperatures, is therefore commonly used as a proxy for cold tolerance (David et al., 1998; Gibert et al., 2001). More severe or prolonged exposure to cold can lead to mortality, which can be quantified either as mortality following a defined cold exposure (cold shock mortality) or as the duration of exposure resulting in 50% mortality (LTi50) (Andersen et al., 2015). Although these phenotypes appear linked, the physiological mechanisms underlying them differ (reviewed in Overgaard & MacMillan, 2017), and different measures of cold tolerance do not necessarily correlate.

Studies of insect cold tolerance have frequently focused on *Drosophila* species, which provide well-established model systems for investigating the physiological and genetic basis of thermal adaptation (Andersen et al., 2015; Bubliy et al., 2002; Colinet et al., 2017; Hoffmann et al., 2002; Hoffmann & Watson, 1993; MacMillan et al., 2015, 2016, 2017; Parker et al., 2015, 2021). Among these, *Drosophila ananassae*, a species of tropical origin, has recently emerged as a promising model for studying the genetic architecture of cold tolerance (Königer & Grath, 2018; Königer et al., 2019; Yılmaz et al., 2023). This species has expanded its range and shows variation in cold tolerance both within and among populations (Tobari, 1993; Das et al., 2004; Königer & Grath, 2018; Königer et al., 2019; Yılmaz et al., 2025). In particular, an ancestral population from Bangkok, Thailand, shows substantial variation among iso-female lines, ranging from fast-recovering, cold-tolerant lines to slow-recovering, cold-sensitive lines. In contrast, a derived population from Kathmandu, Nepal, shows less variation in CCRT, with most iso-female lines characterized as cold tolerant. These patterns make *D. ananassae* a useful system for investigating the genetic basis of variation in cold tolerance.

Previous studies of cold tolerance in *D. ananassae* have primarily focused on CCRT as the phenotypic measure. A comparative transcriptomic study during recovery from chill coma identified genes associated with differences between fast- and slow-recovering iso-female lines from the ancestral Bangkok population (Königer & Grath, 2018). In a subsequent study, recombinant inbred lines (RILs) were generated from the fastest and slowest recovering iso-female lines of this population, and quantitative trait locus (QTL) mapping identified loci that together explained approximately 64% of the phenotypic variance in CCRT (Königer et al., 2019). More recently, transcriptomic analyses of cold acclimation responses revealed biochemical pathways associated with improved CCRT in both Bangkok and Kathmandu populations (Yılmaz et al., 2025). However, these studies focused on a single proxy trait, leaving open whether other measures provide complementary information about cold tolerance.

Bulk segregant analysis (BSA) provides an efficient approach for identifying genomic regions associated with phenotypic variation by comparing allele frequencies between pools of individuals with extreme phenotypes (Ehrenreich et al., 2010; Pool, 2016; Kurlovs et al., 2019). By sequencing pools of individuals from recombinant populations, loci contributing to trait variation can be detected as shifts in allele frequencies between contrasting phenotypic groups. This strategy has been widely used to investigate the genetic architecture of complex traits in *Drosophila* and other organisms, including locomotor activity (Benowitz et al., 2019), pigmentation evolution (Bastide et al., 2016), and zinc toxicity resistance using extreme QTL (X-QTL) mapping (Hanson et al., 2025). Similar pooled-sequencing approaches have also been applied to dissect complex traits in other insects, yeast, and plants (Edwards & Gifford, 2012; Kurlovs et al., 2019; Ma et al., 2023; Magwene et al., 2011; Michelmore et al., 1991; Vos et al., 2022).

Here, we further characterized cold tolerance in iso-female lines from the Bangkok and Kathmandu populations and in 16 RILs derived from the Bangkok population by measuring cold shock (CS) mortality and LTi50 in addition to CCRT. We tested the relationships among these cold tolerance phenotypes and performed BSA using RILs with extreme CCRT phenotypes to identify genomic regions associated with variation in chill coma recovery. Finally, we performed gene ontology enrichment analyses and identified candidate genes within the putative regions. Together, our results emphasize the importance of considering multiple phenotypes when characterizing cold tolerance and provide further insight into its genetic architecture in *D. ananassae*.

## MATERIALS AND METHODS

### Fly Husbandry

This study used eight *Drosophila ananassae* iso-female lines from Thailand (Bangkok, BKK), three iso-female lines from Nepal (Kathmandu, KATH), and sixteen recombinant inbred lines (RILs) previously generated by Königer et al. (2019). The RIL population was established from two Bangkok founder strains with contrasting chill coma recovery times (fast: BKK12; slow: BKK13). Reciprocal crosses (BKK12 males × BKK13 females and BKK13 males × BKK12 females) were performed, and F1 offspring from both crosses were combined and maintained in mass breeding until the F4 generation to promote recombination. From the F5 generation onwards, independent inbred lines were established by full-sib mating, with brother–sister pairs maintained through generation F10. Parents were removed after mating to prevent backcrossing. All iso-female lines were originally collected in 2002 (Das et al., 2004). Fly strains were maintained under standard laboratory conditions (rearing temperature 22 ± 1 °C, 14:10 h light:dark cycle) at low density on corn–molasses medium. All experiments were performed on flies expanded for two generations under identical laboratory conditions.

### Cold Tolerance Phenotype Measurements

For all phenotype measurements, F2 generation flies were collected within 0–2 days of emergence. After one day of recovery, flies were briefly anesthetized with CO₂, separated by sex, and placed into small vials containing fresh fly food in groups of ten individuals. Flies were allowed to recover for an additional day after sex separation and were tested at 4–6 days of age. Three replicate vials were used per sex and strain for all phenotyping assays.

#### Chill Coma Recovery Time

To induce chill coma, 4–6-day-old flies were transferred into empty vials without CO₂ exposure and placed on melting ice at 0 °C for 3 hours. After cold exposure, flies were returned to the rearing temperature, and the time required for recovery (defined as regaining the ability to stand) was recorded for up to 90 minutes. Flies that had not recovered after 90 minutes were assigned a value of 92 minutes, and dead flies were excluded from the analysis.

#### Cold Shock Mortality and LTi50

Four- to six-day-old flies were transferred into empty vials without CO₂ exposure and placed on melting ice at 0 °C for exposure periods ranging from 1 to 24 hours, spanning approximately 0–100% mortality. After cold exposure, flies were returned to the rearing temperature for 2 hours to allow recovery from chill coma, after which the number of dead flies in each vial was recorded. Mortality after 8 hours of exposure was used to calculate cold shock mortality. LTi50 values were determined as the exposure duration corresponding to 50% mortality for each sex and strain based on the full mortality dataset.

#### Data Analysis

All statistical analyses were performed in R version 4.3.3 (R Core Team, 2022). LTi50 values were estimated using generalized linear models with a quasibinomial error distribution, and 95% confidence intervals were calculated using the Delta method (Oehlert, 1992). The effects of sex and strain on chill coma recovery time (CCRT) and cold shock mortality were assessed using two-way analyses of variance (ANOVA), including the interaction between sex and strain (`Phenotype ∼ Sex * Strain`). For LTi50, an additive two-way ANOVA model (`LTi50 ∼ Sex + Strain`) was used because only one LTi50 estimate was available per sex–strain combination, preventing estimation of the interaction effect. Statistical significance was assessed at α = 0.05.

Relationships among chill coma recovery time (CCRT), cold shock mortality, and LTi50 were assessed using Spearman’s rank correlation coefficient. To evaluate whether the inclusion of founder and population lines influenced the observed associations, correlations were calculated for both the complete dataset and for the recombinant inbred lines (RILs) only, as the latter represent the genetically segregating mapping population. Correlations were calculated using mean phenotypic values for each sex and strain. Statistical significance was assessed at α = 0.05.

### Sample Collection and DNA Extraction

Four pools were generated for bulk segregant analysis. For the parental pools, 100 females were collected from the F2 generations of each founder strain. For recombinant inbred line pools with extreme phenotypes, flies from the F2 generation of selected RILs (line numbers 22, 23, 57, 80, 90, 93) were collected within 0–2 days of emergence. Selection of extreme phenotypes was based solely on chill coma recovery time (CCRT). After one day of recovery, flies were briefly anesthetized with CO₂ and separated by sex. Individual females were then placed into small vials containing fresh fly food.

At 4–6 days of age, flies were placed on melting ice at 0 °C for 3 hours to induce chill coma. The vials were subsequently returned to the rearing temperature for recovery and checked every 30 minutes. Flies that regained the ability to stand within the first 30 minutes were collected as extremely fast-recovering individuals. Flies that recovered during the second 30-minute interval were discarded. After the third and final 30-minute interval, all remaining living flies were collected as extremely slow-recovering individuals regardless of their recovery state. This procedure was repeated until 100 flies were obtained for each extreme pool. Extreme tolerant and sensitive bulks were constructed by pooling individuals from RILs with the lowest and highest CCRT values, respectively.

Flies were placed into 1.5 mL Eppendorf tubes in groups of ten and snap-frozen in liquid nitrogen prior to DNA extraction. Genomic DNA was extracted using the MasterPure™ DNA Purification Kit (Epicentre, Madison, WI, USA). DNA concentration and purity were assessed using both a NanoDrop spectrophotometer (Thermo Fisher Scientific) and a Qubit fluorometer (Thermo Fisher Scientific).

### Whole Genome Sequencing

DNA samples were pooled and submitted for whole genome sequencing to an external facility (Novogene, Cambridge, UK). High-throughput sequencing was performed on a NovaSeq 6000 platform (Illumina, San Diego, CA, USA) generating 150-bp paired-end reads. On average, 62.4 million reads were generated per sample (Table S5). Adapter trimming and quality filtering were performed by the sequencing provider.

Raw reads were mapped to the *Drosophila ananassae* reference genome (NCBI *D. ananassae* Annotation Release 102, accession ASM1763931v2) using the Burrows–Wheeler Aligner (BWA; version 0.5.0). Subsequent processing steps, including sorting, merging, and duplicate removal, were performed using SAMtools (version 1.18). Variant calling was conducted using FreeBayes (version 1.3.8), and variants were filtered using BCFtools (version 1.18). Finally, the GATK tool VariantsToTable was used to convert variant data into tabular format for downstream analysis in R (version 4.2.2).

### Bulk Segregant Analysis

Bulk segregant analysis (BSA) was performed to identify genomic regions associated with variation in chill coma recovery time (CCRT). In this approach, individuals with extreme phenotypes are pooled and sequenced, and differences in allele frequencies between pools are used to detect loci linked to the trait of interest (Pool, 2016; Kurlovs et al., 2019).

The analysis was restricted to both arms of chromosomes 2 and 3 and the X chromosome, while unlocalized scaffolds were excluded. Individuals from recombinant inbred lines (RILs) exhibiting extreme CCRT phenotypes were assigned to two pools (100 individuals per pool) representing tolerant (low CCRT) and sensitive (high CCRT) flies. Genomic regions associated with phenotypic differences between the two pools were identified using the QTLseqr package (version 0.7.5.2) (Mansfeld & Grumet 2018) in R.

For each SNP, the SNP-index was calculated for both pools, and the Δ(SNP-index) was determined as the difference in alternate allele frequencies between the tolerant and sensitive bulks. SNPs were filtered based on sequencing depth and quality prior to analysis (parameters QUAL ≥ 10; DP > 100). In addition, G statistics for individual SNPs and smoothed G’ values within 1 Mb sliding windows were calculated. While Δ(SNP-index) highlights SNPs with large allele frequency differences between bulks, the G’ statistic detects regions showing consistent allele frequency divergence across multiple SNPs. Genomic regions exceeding the significance thresholds of these analyses were considered candidate QTLs and were subsequently annotated using the gene annotation file associated with the reference genome.

### Gene Ontology Enrichment Analysis

Genes located within the candidate regions were identified. Gene ontology (GO) enrichment analysis was performed on the *D. melanogaster* orthologs of these genes using the R package clusterProfiler (version 4.6.2) (Wu et al., 2021). P-values were adjusted for multiple testing using the false discovery rate (FDR) method with a significance threshold of 0.01. Significant GO terms were visualized using the R package genekitr (version 1.2.5) (Liu & Li, 2023).

### Ethics statement

No ethical approval or specific permissions were required for this study, as it involved work with *Drosophila ananassae*, an invertebrate species not subject to institutional or national animal research regulations.

## RESULTS

To characterize variation in cold tolerance, we measured three phenotypes, chill coma recovery time (CCRT), cold shock mortality, and lethal time (LTi50), in iso-female lines from the Bangkok and Kathmandu populations and in recombinant inbred lines (RILs).

### Variation in Chill Coma Recovery Time among Populations and RILs

Chill coma recovery time (CCRT) varied substantially among iso-female lines and recombinant inbred lines (RILs) of *Drosophila ananassae* following a 3 h cold exposure (Figure 1A, Table S1, Table S9). Mean CCRT across both sexes was 42.5 minutes for fast-recovering Bangkok lines, 61.0 minutes for slow-recovering Bangkok lines, and 55.3 minutes for Kathmandu lines. In the RIL panel, average CCRT was 54.62 minutes and ranged from 28.0 to 84.3 minutes.

**Figure 1.**
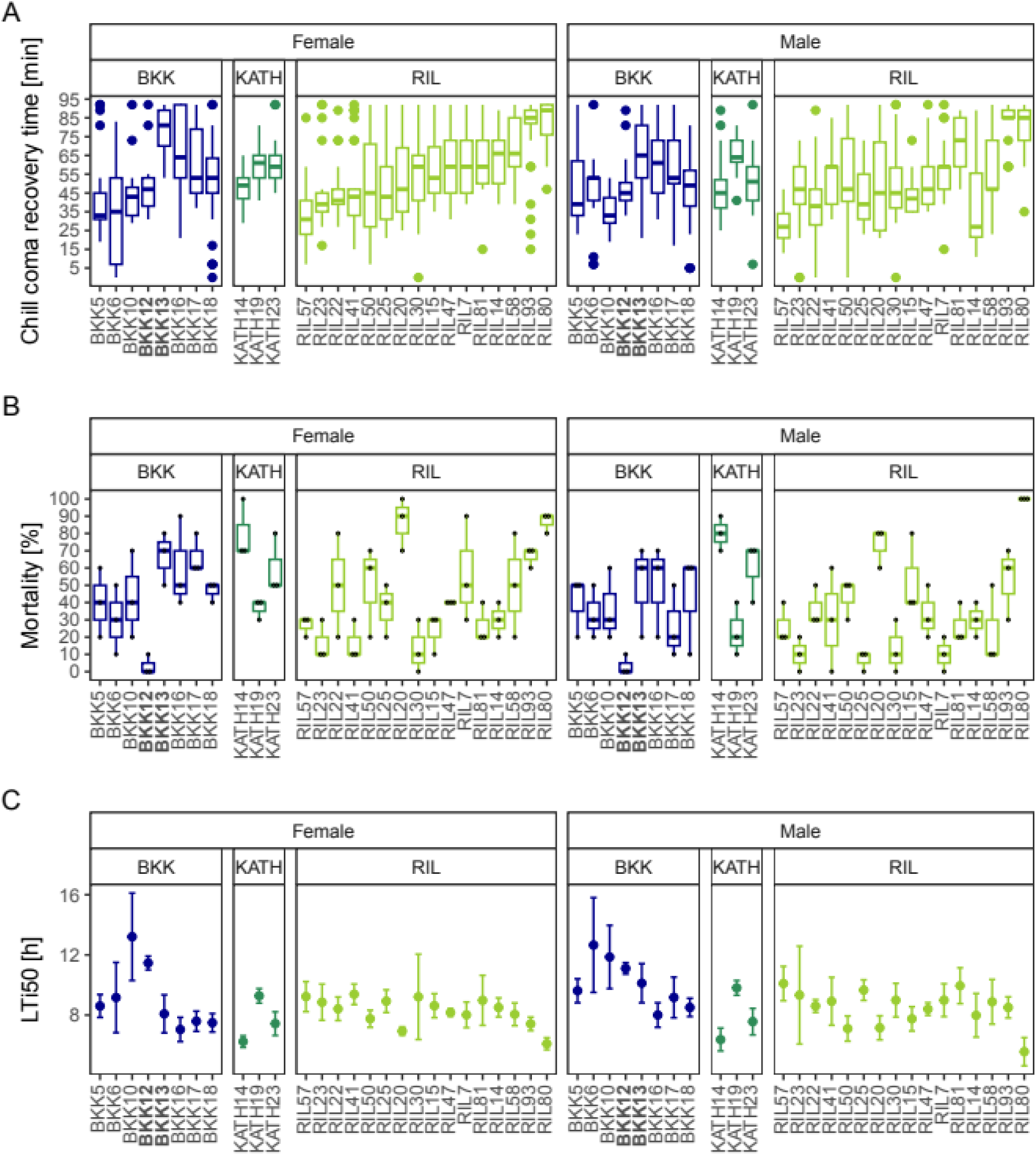
Cold tolerance phenotypes in *Drosophila ananassae* iso-female lines and recombinant inbred lines (RILs). (A) Chill coma recovery time (CCRT) measured after a 3 h cold shock. Thirty flies per sex and strain were tested in groups of ten. (B) Mortality (%) measured after an 8 h cold shock followed by 2 h recovery. For each strain and sex, three replicate vials containing ten flies each were tested; large colored dots indicate replicate values. For panels A and B, boxes indicate the interquartile range, horizontal lines represent the median, whiskers extend to 1.5× the interquartile range, and outliers are shown as colored dots. (C) Lethal time to 50% mortality (LTi50) in hours. LTi50 was estimated using generalized linear models with a quasibinomial distribution; dots indicate estimated values for each strain and sex, and error bars represent 95% confidence intervals. Female phenotypes are shown in the left panels and male phenotypes in the right panels. Colors indicate the origin of the lines: Bangkok (dark blue, BKK), Kathmandu (dark green, KATH), and recombinant inbred lines (light green, RIL). Founder strains used to generate the RIL population are indicated in bold. RILs are ordered in ascending order according to female CCRT values. *Alt text: Boxplots and dot plots comparing cold tolerance phenotypes across populations and recombinant inbred lines. Chill coma recovery time and mortality vary among strains, while lethal time estimates show substantial variation, with some recombinant lines exceeding founder ranges*.

A two-way ANOVA revealed significant effects of sex, strain, and a significant Sex × Strain interaction on chill coma recovery time (Table 1). Sex-specific values for all three cold-tolerance phenotypes are shown in Figure S1, with female and male values connected within strains to visualize the magnitude and direction of sex differences.

**Table 1:** Results of a two-way ANOVA for chill coma recovery time (CCRT) using the model CCRT ∼ Sex × Strain. The interaction term (Sex x Strain) is included in the model. Degrees of freedom (df), sums of squares (Sum Sq), mean squares (Mean Sq), *F* values and corresponding *P* values are shown.

|  | df | Sum Sq | Mean Sq | F value | P value |
| --- | --- | --- | --- | --- | --- |
| Sex | 1 | 3,946 | 3,946 | 13.043 | 0.000314 |
| Strain | 26 | 223,470 | 8,595 | 28.411 | $< 2 \times 10^{-16}$ |
| Sex:Strain | 26 | 28,167 | 1,083 | 3.581 | $3.68 \times 10^{-9}$ |
| Residuals | 1548 | 468,307 | 303 |  |  |

Several RILs exhibited recovery times beyond the range of the founder strains, consistent with transgressive phenotypes. RILs with recovery times below the observed value of the fast founder (BKK12) or above that of the slow founder (BKK13) were classified as extreme phenotypes. The three RILs with the lowest and the three RILs with the highest CCRT values were selected for bulk segregant analysis. Individuals from the three lines in each group were pooled to generate the tolerant and sensitive sequencing bulks.

### Cold Shock Mortality Differs among Strains and Sexes

Mortality following cold shock was quantified after 8 h of cold exposure followed by 2 h of recovery (Figure 1B, Table S2, Table S10). Mean mortality was 28.7% for fast-recovering Bangkok strains, 51.3% for slow-recovering Bangkok strains, and 56.7% for Kathmandu strains. Among the RILs, mean mortality was 39.0%, ranging from 10.0% to 86.7%.

Two-way ANOVA indicated significant effects of sex and strain on mortality but no significant interaction between these factors (Table 2).

**Table 2:** Results of a two-way ANOVA for cold shock mortality using the model Mortality ∼ Sex × Strain. The interaction term (Sex x Strain) is included in the model. Degrees of freedom (df), sums of squares (Sum Sq), mean squares (Mean Sq), *F* values and corresponding *P* values are shown.

|  | df | Sum Sq | Mean Sq | F value | P value |
| --- | --- | --- | --- | --- | --- |
| Sex | 1 | 20.8 | 20.765 | 6.908 | 0.00983 |
| Strain | 26 | 693.9 | 26.687 | 8.877 | $< 2 \times 10^{-16}$ |
| Sex:Strain | 26 | 98.2 | 3.778 | 1.257 | 0.20706 |
| Residuals | 108 | 324.7 | 3.006 |  |  |

### Lethal Time (LTi50) Varies among Strains and Sexes

The third phenotype examined was lethal time (LTi50), defined as the duration of cold exposure resulting in 50% mortality. Mortality rates measured across multiple exposure durations were used to estimate LTi50 values (Figure 1C, Table S3).

The average LTi50 was 10.9 h for fast-recovering Bangkok lines, 8.3 h for slow-recovering Bangkok lines, and 7.8 h for Kathmandu strains. Among the RILs, LTi50 values ranged from 5.6 to 10.1 h.

Because LTi50 was estimated once for each sex and strain, the interaction between sex and strain could not be tested. We therefore used an additive two-way ANOVA including sex and strain as main effects. Both strain and sex had significant effects on LTi50 (Table 3), with males showing an approximately 0.46 h (28 min) higher LTi50 than females.

**Table 3:** Results of an additive two-way ANOVA for lethal time (LTi50) using the model LTi50 ∼ Sex + Strain. The additive model includes the main effects of sex and strain but no interaction term, because only one LTi50 estimate was available for each sex–strain combination. Degrees of freedom (df), sums of squares (Sum Sq), mean squares (Mean Sq), F values, and corresponding P values are shown.

|  | df | Sum Sq | Mean Sq | F value | P value |
| --- | --- | --- | --- | --- | --- |
| Sex | 1 | 2.91 | 2.907 | 5.861 | 0.0228 |
| Strain | 26 | 105.58 | 4.061 | 8.185 | $4.39 \times 10^{-7}$ |
| Residuals | 26 | 12.90 | 0.496 |  |  |

### Cold Tolerance Phenotypes Capture Distinct Components of the Cold Response

To assess relationships among the three cold tolerance phenotypes, we calculated Spearman rank correlations using both the complete dataset and the recombinant inbred lines (RILs) only (Table 4). Correlation coefficients were similar between both datasets. In both analyses, cold shock mortality showed a strong negative correlation with LTi50 (ρ = –0.77 and ρ = –0.86 for all lines and RILs, respectively), whereas CCRT showed weaker correlations with mortality (ρ = 0.34 and ρ = 0.36) and LTi50 (ρ = –0.33 in both datasets). Overall, these results indicate that the three measures of cold tolerance capture partially overlapping but distinct aspects of the cold response.

**Table 4:** Spearman rank correlations among cold tolerance phenotypes calculated for all lines and for the recombinant inbred lines (RILs) only. Correlation coefficients (*ρ*), associated P-values, and sample sizes (n) are shown.

| Phenotype 1 | Phenotype 2 | Lines | Spearman's $\rho$ | P value | n |
| --- | --- | --- | --- | --- | --- |
| CCRT | Mortality | RILs | 0.357826 | 0.04435 | 32 |
| CCRT | Mortality | All | 0.3363177 | 0.0129 | 54 |
| CCRT | LTi50 | RILs | -0.3302786 | 0.06538 | 32 |
| CCRT | LTi50 | All | -0.3313512 | 0.01476 | 54 |
| Mortality | LTi50 | RILs | -0.8602152 | $2.792 \times 10^{-10}$ | 32 |
| Mortality | LTi50 | All | -0.7664684 | $1.424 \times 10^{-11}$ | 54 |

### Bulk Segregant Analysis Identifies Candidate Genomic Regions for CCRT

To identify genomic regions associated with variation in chill coma recovery time, we performed bulk segregant analysis (BSA) using individuals from recombinant inbred lines with extreme CCRT phenotypes together with the founder strains. Each sequencing pool comprised 100 female flies. After quality filtering, 301,000 SNPs remained for analysis (Figure 2A, Table S6). Genomic differentiation between tolerant and sensitive pools was assessed using two complementary approaches: the Δ(SNP-index), which identifies allele frequency differences between pools (Takagi et al., 2013), and the G′ statistic, which incorporates information from neighbouring SNPs (Magwene et al., 2011). The Δ(SNP-index) analysis identified 22 candidate genomic regions associated with variation in CCRT (Figure 2B, Table S7). Ten regions showed negative Δ(SNP-index) values, whereas twelve regions showed positive values. G′ peaks broadly overlapped with regions identified by the Δ(SNP-index) analysis; however, no regions remained significant after correction for multiple testing (Figure S2). We therefore consider the Δ(SNP-index) regions as candidate loci requiring further validation.

**Figure 2.**
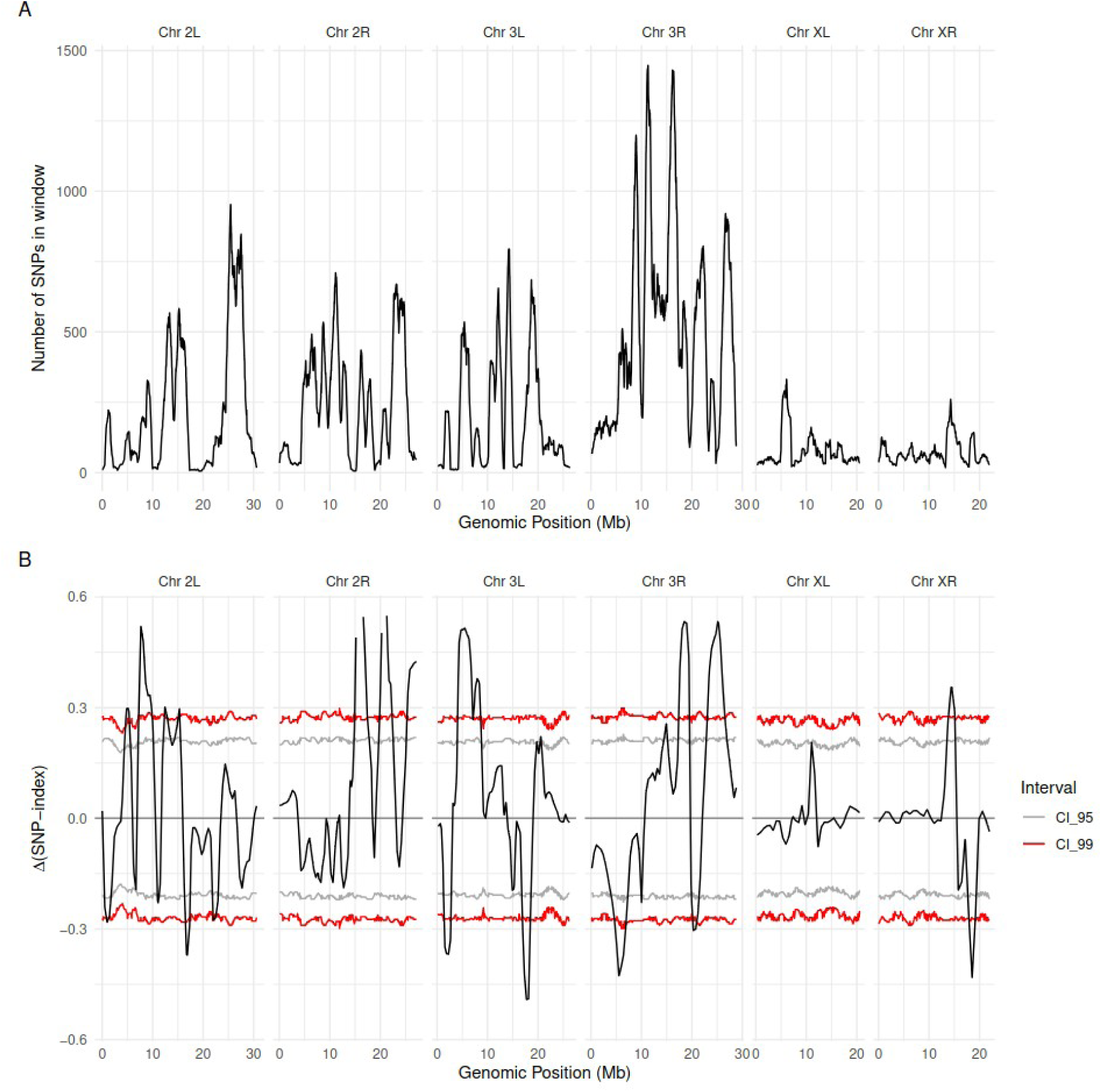
QTL-seq analysis of variation in chill coma recovery time. (A) SNP density across the genome calculated within 1 Mb windows. (B) Δ(SNP-index) values across the genome comparing pools of extremely tolerant and extremely sensitive recombinant inbred lines. Shaded regions indicate the 95% confidence interval (gray) and 99% confidence interval (red). Genomic position is shown on the x-axis, with chromosomes indicated above the panels. QTL regions exceeding the 99% confidence interval were considered significantly associated with the phenotype. *Alt text: Genome-wide plots showing SNP density and allele frequency differences between tolerant and sensitive pools. Multiple genomic regions exceed confidence thresholds, indicating candidate loci associated with variation in chill coma recovery time*.

### Candidate Regions are Enriched for Genes Involved in Muscle and Metabolic Processes

Within the candidate regions identified by the Δ(SNP-index) analysis, a total of 2,379 genes were located. To investigate the potential biological functions represented within these regions, we performed gene ontology (GO) enrichment analysis using the *D. melanogaster* orthologs of these genes.

The analysis revealed enrichment of GO terms related to muscle development and metabolic processes (Figure 3, Table S8). When the analysis was separated according to Δ(SNP-index) direction, regions with positive Δ(SNP-index) values were enriched for GO terms associated with muscle development, metabolic processes, and cytoskeletal protein binding (Figure S3), whereas regions with negative Δ(SNP-index) values were enriched for glucosidase, monooxygenase, and hydrolase activities (Figure S4, Table S8).

**Figure 3.**
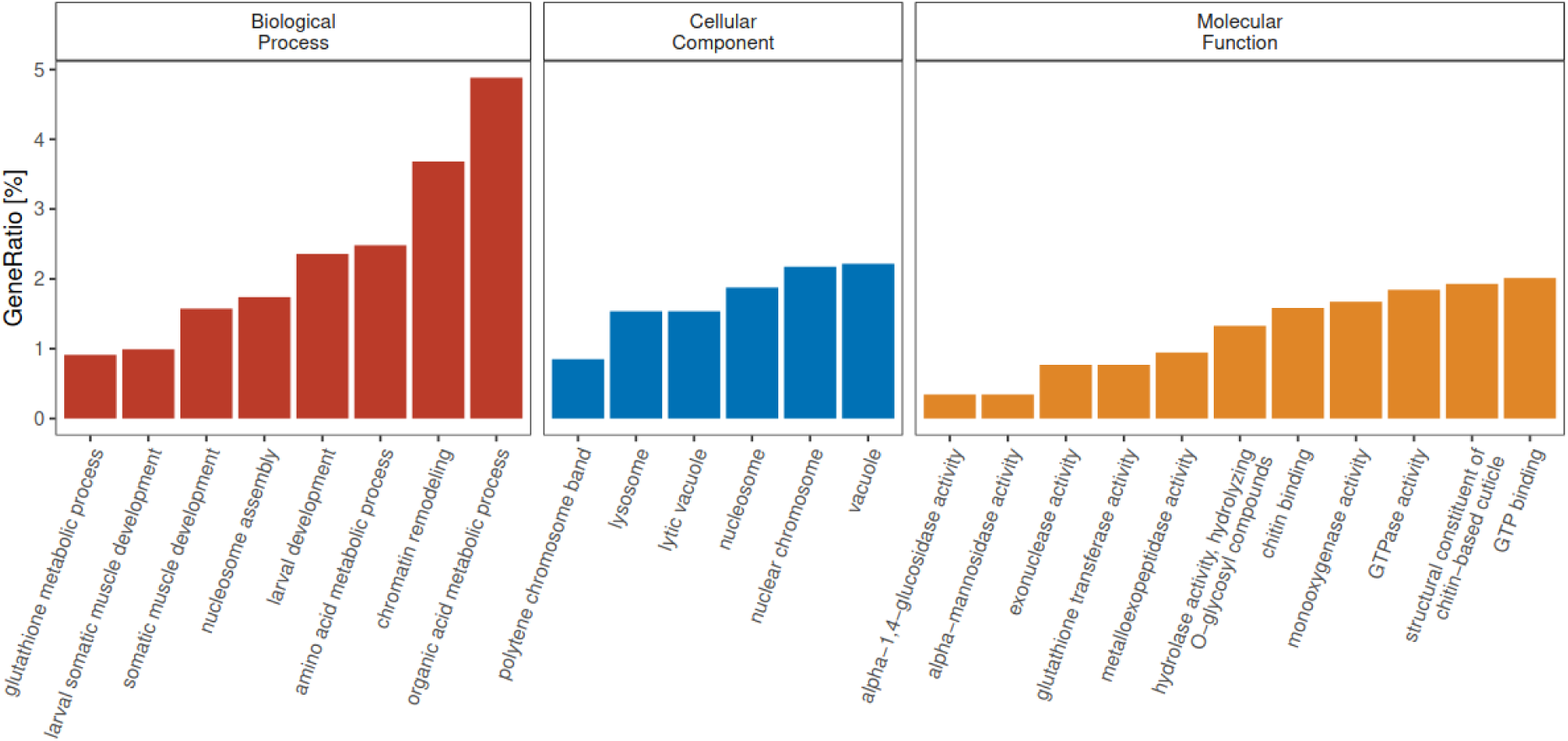
Gene ontology (GO) terms enriched in genes located within candidate regions. The top 25 GO terms ranked by adjusted p-value are shown. The y-axis indicates the ratio of genes associated with each GO term. Biological processes are shown in red, cellular components in blue, and molecular functions in yellow. *Alt text: Bar plots of gene ontology enrichment for genes in candidate regions, highlighting overrepresentation of functions related to muscle development, cytoskeletal organization, and metabolic processes*.

Taken together, these analyses reveal substantial variation in cold tolerance among populations and recombinant inbred lines of *Drosophila ananassae*. The three phenotypic assays showed partly overlapping but distinct patterns of variation, and recombinant lines frequently exhibited extreme phenotypes relative to the founder strains. In addition, bulk segregant analysis identified multiple genomic regions associated with variation in chill coma recovery time, with candidate genes enriched for functions related to muscle development and metabolic processes.

## DISCUSSION

The three phenotypes measured here capture partially overlapping but distinct aspects of the cold response. Together, these results indicate that cold tolerance in *Drosophila ananassae* is a complex trait composed of partially independent phenotypic components. In particular, we observed substantial variation among iso-female lines and between sexes, and the different measures of cold tolerance did not always show consistent patterns across populations. Consistent with this interpretation, some recombinant inbred lines consistently exhibited extreme phenotypes across multiple assays (e.g., RIL80 and RIL57), whereas others were extreme only for specific traits.

Our phenotypic comparisons among Bangkok and Kathmandu iso-female lines further highlight the complexity of cold tolerance variation within and between populations. Previous analyses suggested that four iso-female lines of the Bangkok (BKK) population were slow-recovering with high CCRT, whereas four lines from the same population and three lines from the Kathmandu (KATH) population were fast-recovering with low CCRT (Königer & Grath, 2018). While our measurements confirmed that the fast-recovering BKK lines showed lower average CCRT than both the slow-recovering BKK and KATH lines, additional phenotypic characterization revealed more complex and trait-specific patterns. For example, males of the slow-recovering BKK lines were more tolerant than males of the KATH lines based on CS mortality and LTi50, whereas females showed the opposite pattern. We also observed considerable variation among iso-female lines within each group. Together, these results highlight the importance of using multiple phenotypic measures when characterizing cold tolerance.

Recombination among founder genotypes revealed extensive phenotypic variation in the recombinant inbred lines. Recombinant inbred lines showed continua of CCRT, LTi50, and CS mortality. For CCRT, some RILs exhibited extreme phenotypes at both ends of the distribution. In other words, some RILs recovered from chill coma faster than the fast-recovering founder BKK12, whereas others recovered more slowly than the slow-recovering founder BKK13. In contrast, extreme phenotypes for LTi50 and CS mortality were observed only at the sensitive end, with lower LTi50 values and higher mortality than BKK13. Importantly, CCRT was used to determine the founder strains, while LTi50 and CS mortality were not measured in the iso-female lines prior to generating the RILs.

The continuum of phenotypes and the presence of transgressive values indicate that cold tolerance is influenced by multiple genetic factors segregating within the ancestral population. Recombination among founder genotypes can generate novel combinations of alleles that produce phenotypes outside the range of the original strains. Such patterns are consistent with a complex, polygenic trait architecture and demonstrate the presence of standing genetic variation affecting cold tolerance within the Bangkok population. These patterns are consistent with transgressive segregation arising from recombination among alleles from the founder strains (Rieseberg et al., 1999).

Our measurements also revealed differences between current CCRT estimates and previously reported values for the RIL population. The CCRT measurements of the RILs differed from those reported by Königer et al. (2019), as the line rankings based on initial CCRT values did not align with the averages measured here. Such changes in phenotype may reflect residual genetic variation within lines or shifts in allele frequencies due to bottlenecks or inbreeding during line maintenance. We therefore selected individuals from RILs exhibiting extremely low or high CCRT based on the current phenotypic measurements for bulk segregant analysis.

Using bulk segregant analysis, we identified 22 candidate genomic regions associated with variation in CCRT. Comparison of Δ(SNP-index) and G′ analyses revealed partially overlapping signals, but only the Δ(SNP-index) approach detected regions exceeding significance thresholds. This difference likely reflects the distinct statistical properties of the two methods: Δ(SNP-index) is sensitive to localized allele frequency differences, whereas the G′ statistic captures broader, consistent signals across genomic regions (Magwene et al., 2011; Takagi et al., 2013). The lack of significance in the G′ analysis therefore does not necessarily contradict the Δ(SNP-index) results, but instead suggests that some signals may be driven by relatively narrow genomic intervals. Accordingly, the regions identified here should be considered putative loci requiring further validation.

One of the identified candidate regions contains genes potentially involved in processes relevant to cold tolerance. SNPs in this region exhibited *p*-values below 0.05 prior to multiple-testing correction, making it a candidate for further validation. This region spans a 682-kb interval on chromosome 2R and contains genes enriched for functions related to muscle development, chitin binding, and palmitoylation (Table S4). Among the 19 chitin-binding genes in this region, the *D. melanogaster* orthologs of GF25314 and GF24516 (CG17147 and Muc68D, respectively) were previously associated with cold acclimation (MacMillan et al., 2016). Two additional orthologs, GF24478 (CG6933) and GF24481 (CG7298), show high or very high expression in response to cold, although they have not previously been directly linked to cold tolerance. Previous work also suggested that chitin-binding genes play an important role in the cold acclimation response of *D. ananassae* (Yılmaz et al., 2025), where their expression increased in both tolerant and sensitive strains during acclimation. The present results are consistent with a role of chitin-binding genes in cold tolerance.

Several genes within the candidate regions are also associated with muscle development. For example, *polo* has previously been linked to hypoxia tolerance (Azad et al., 2012; Gilliland et al., 2009), while the human ortholog of *rt* is associated with muscular dystrophy (Jurado et al., 1999). In addition, four *D. melanogaster* genes associated with protein palmitoylation in the GO analysis correspond to a single *D. ananassae* ortholog, GF24551, linking protein palmitoylation to one of the candidate regions identified here.

Gene ontology analysis of genes located within the candidate regions highlighted several biological processes potentially involved in chill coma recovery, including muscle development, cytoskeletal organization, chitin binding, and protein palmitoylation. Chill coma recovery requires restoration of membrane potential and ion-water balance, ultimately leading to the recovery of neuromuscular coordination (Overgaard & MacMillan, 2017). As the recovery of muscle control is essential for chill coma recovery, variation in genes involved in muscle development may contribute to differences in CCRT.

Several candidate genes identified in these regions are consistent with these processes. For example, *Obscurin* and *Msp300* encode proteins involved in sarcomere assembly and the positioning of muscle nuclei and mitochondria, respectively (Volk, 1992; Schnorrer et al., 2010). Neither of these two genes has previously been associated with cold-tolerance phenotypes. Cytoskeletal elements have previously been implicated in cold stress responses in insects, and genes associated with actin dynamics show differential expression during both cold shock and cold acclimation (Königer & Grath, 2018; Yılmaz et al., 2025). The cytoskeleton plays an important role in maintaining cellular structure and anchoring ion channels, which may influence ion homeostasis during cold stress (Denker & Barber, 2002). Consistent with this, the cytoskeleton-related gene *klarsicht* was previously suggested as a candidate gene underlying differences between fast- and slow-recovering BKK strains (Königer et al., 2019). Like *Msp300*, *klarsicht* is involved in positioning nuclei in muscle. Two additional genes located within the candidate regions, GF14829 (*D. melanogaster* CG10383) and GF21827 (*D. melanogaster Arpc3B*), have been linked to adult longevity and sleep, and actin binding, respectively (Paik et al., 2012; Hudson & Cooley, 2002), and were previously associated with phenotypic differences within the BKK population (Königer & Grath, 2018; Königer et al., 2019).

In addition to cytoskeletal and muscle-related genes, we observed enrichment for genes involved in chitin binding and protein palmitoylation. Palmitoylation is the covalent attachment of palmitic acid, a 16-carbon fatty acid, to cysteine residues of proteins by thioester bonds (Linder & Deschenes, 2007). It regulates protein localization, protein–protein interactions, signal transduction, protein stability, and membrane organization. Nine *D. ananassae* genes within the candidate regions showed enrichment in protein palmitoylation, four of which have previously been shown to be expressed in the central nervous system (Bannan et al., 2008). Because membrane fluidity and cellular signaling are strongly affected by low temperatures, variation in palmitoylation-related genes may contribute to differences in cold tolerance.

Exposure to low temperatures disrupts cellular homeostasis by affecting protein stability, energy metabolism, and membrane fluidity (Overgaard & MacMillan, 2017; Teets & Denlinger, 2013). Extended exposure to cold leads to chill injury and, eventually, insect death. Maintaining ion-water balance is considered a key determinant of cold tolerance in chill-susceptible insects (Andersen et al., 2017; Overgaard et al., 2021). Disturbance of ion homeostasis leads to loss of membrane potential and impaired muscle excitability (Overgaard & MacMillan, 2017). However, it remains uncertain whether the loss of neuromuscular coordination results primarily from failure in neurons, muscles, or synapses. Our results suggest that, in addition to processes previously associated with cold tolerance, including ion transport, fatty acid metabolism, protein folding, and cellular respiration, genes involved in muscle development and palmitoylation may contribute to variation in cold tolerance and provide further evidence for roles of the cytoskeleton and chitin binding in chill coma recovery.

Identifying candidate loci associated with cold tolerance provides a foundation for functional validation. The availability of genome editing tools for *D. ananassae* (Yılmaz et al., 2023) provides opportunities to experimentally validate the candidate genes identified in this study. In particular, 16 candidate genes (Table S4) could be investigated using genome editing followed by phenotype evaluation, electrophysiological assays, and reabsorption measurements. Such functional studies could clarify the roles of individual candidate genes in chill coma recovery and cold tolerance. In addition, selection experiments using strains with contrasting genotypes may help reveal how selection acts on the genetic variation underlying these traits. Overall, by integrating multiple phenotypic measurements with genome-wide variant analysis, our study provides new insights into the genetic basis of chill coma recovery and cold tolerance in *Drosophila ananassae*.

## AUTHOR CONTRIBUTIONS

Conceptualization: S.G., V.M.Y.; Methodology: S.G., V.M.Y.; Software: V.M.Y., F.T.K.; Validation: S.G., V.M.Y., V.O.; Formal analysis: V.M.Y., F.T.K.; Investigation: V.M.Y., F.T.K.; Resources: S.G.; Data curation: V.M.Y., F.T.K.; Writing – original draft: V.M.Y.; Writing – review and editing: S.G., V.M.Y., V.O.; Visualization: V.M.Y., V.O.; Supervision: S.G.; Project administration: S.G.; Funding acquisition: S.G.

## ACKNOWLEDGEMENTS

We thank Hilde Lainer and Bianca Hölldobler for technical assistance in the laboratory, Amanda Glaser-Schmitt and John Parsch for their valuable input on experimental design, and Dirk Metzler for help regarding statistical analyses. We extend our appreciation to all current and previous members of the Division of Evolutionary Biology, as well as the Graduate School Life Science Munich (LSM) at LMU Munich. This work was supported by the Deutsche Forschungsgemeinschaft (DFG) (project number 271330745, awarded to S.G.).

## DATA AVAILABILITY

Whole-genome sequencing data have been deposited in the NCBI Sequence Read Archive (SRA) under accession number PRJNA1247020. Analysis scripts and additional data will be made publicly available in a GitHub repository and archived on Zenodo upon acceptance of the manuscript at the time of publication. All other data supporting the findings of this study are included in the article and its Supplementary Material.

## CONFLICTS OF INTEREST

The authors declare no conflict of interest. The funders had no role in the design of the study, in the collection, analyses, or interpretation of data, in the writing of the manuscript, or in the decision to publish the results.

## Supplementary Material

## Supplementary Tables

**Table S1.**
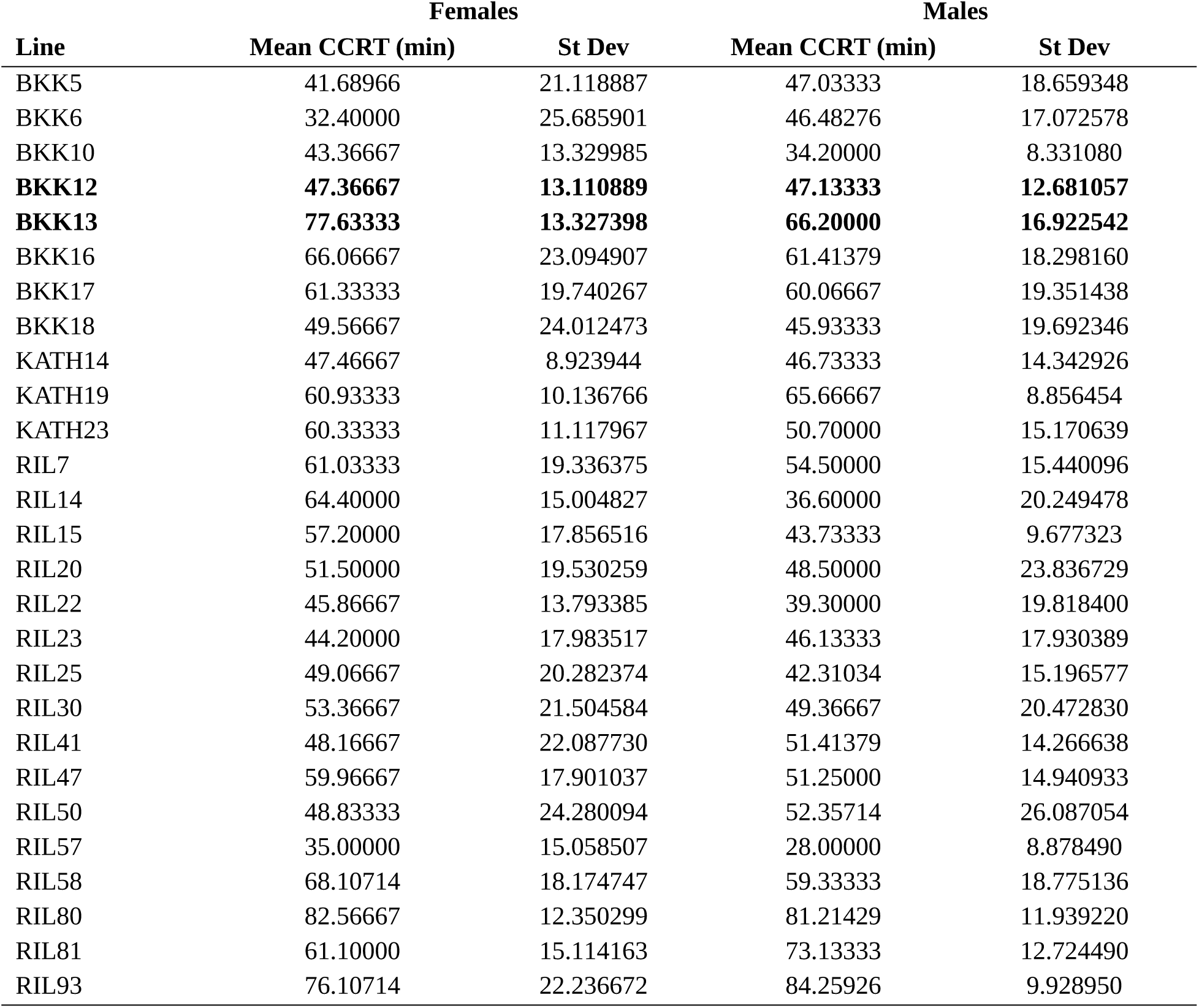
Mean chill coma recovery time (CCRT, minutes) and standard deviation for female and male flies of each strain. RIL founder strains are indicated in **bold**.

| Line | Females |  | Males |  |
| --- | --- | --- | --- | --- |
|  | Mean CCRT (min) | St Dev | Mean CCRT (min) | St Dev |
| BKK5 | 41.68966 | 21.118887 | 47.03333 | 18.659348 |
| BKK6 | 32.40000 | 25.685901 | 46.48276 | 17.072578 |
| BKK10 | 43.36667 | 13.329985 | 34.20000 | 8.331080 |
| **BKK12** | **47.36667** | **13.110889** | **47.13333** | **12.681057** |
| **BKK13** | **77.63333** | **13.327398** | **66.20000** | **16.922542** |
| BKK16 | 66.06667 | 23.094907 | 61.41379 | 18.298160 |
| BKK17 | 61.33333 | 19.740267 | 60.06667 | 19.351438 |
| BKK18 | 49.56667 | 24.012473 | 45.93333 | 19.692346 |
| KATH14 | 47.46667 | 8.923944 | 46.73333 | 14.342926 |
| KATH19 | 60.93333 | 10.136766 | 65.66667 | 8.856454 |
| KATH23 | 60.33333 | 11.117967 | 50.70000 | 15.170639 |
| RIL7 | 61.03333 | 19.336375 | 54.50000 | 15.440096 |
| RIL14 | 64.40000 | 15.004827 | 36.60000 | 20.249478 |
| RIL15 | 57.20000 | 17.856516 | 43.73333 | 9.677323 |
| RIL20 | 51.50000 | 19.530259 | 48.50000 | 23.836729 |
| RIL22 | 45.86667 | 13.793385 | 39.30000 | 19.818400 |
| RIL23 | 44.20000 | 17.983517 | 46.13333 | 17.930389 |
| RIL25 | 49.06667 | 20.282374 | 42.31034 | 15.196577 |
| RIL30 | 53.36667 | 21.504584 | 49.36667 | 20.472830 |
| RIL41 | 48.16667 | 22.087730 | 51.41379 | 14.266638 |
| RIL47 | 59.96667 | 17.901037 | 51.25000 | 14.940933 |
| RIL50 | 48.83333 | 24.280094 | 52.35714 | 26.087054 |
| RIL57 | 35.00000 | 15.058507 | 28.00000 | 8.878490 |
| RIL58 | 68.10714 | 18.174747 | 59.33333 | 18.775136 |
| RIL80 | 82.56667 | 12.350299 | 81.21429 | 11.939220 |
| RIL81 | 61.10000 | 15.114163 | 73.13333 | 12.724490 |
| RIL93 | 76.10714 | 22.236672 | 84.25926 | 9.928950 |

**Table S2.** Mean mortality after an 8 h cold shock and standard deviation for female and male flies of each strain. Mortality represents the number of dead flies per vial (10 flies). RIL founder strains are indicated in **bold**.

| Line | Females |  | Males |  |
| --- | --- | --- | --- | --- |
|  | Mean Mortality | St Dev | Mean Mortality | St Dev |
| BKK5 | 4.00 | 2.0000000 | 4.00 | 1.7320508 |
| BKK6 | 3.00 | 2.0000000 | 3.33 | 1.5275252 |
| BKK10 | 4.33 | 2.5166115 | 3.67 | 2.0816660 |
| **BKK12** | **0.33** | **0.5773503** | **0.33** | **0.5773503** |
| **BKK13** | **6.67** | **1.5275252** | **5.00** | **2.6457513** |
| BKK16 | 6.00 | 2.6457513 | 5.00 | 2.6457513 |
| BKK17 | 6.67 | 1.1547005 | 2.67 | 2.0816660 |
| BKK18 | 4.67 | 0.5773503 | 4.33 | 2.8867513 |
| KATH14 | 8.00 | 1.7320508 | 8.00 | 1.0000000 |
| KATH19 | 3.67 | 0.5773503 | 2.33 | 1.5275252 |
| KATH23 | 6.00 | 1.7320508 | 6.00 | 1.7320508 |
| RIL7 | 5.67 | 3.0550505 | 1.00 | 1.0000000 |
| RIL14 | 3.00 | 1.0000000 | 3.00 | 1.0000000 |
| RIL15 | 2.33 | 1.1547005 | 5.33 | 2.3094011 |
| RIL20 | 8.67 | 1.5275252 | 7.33 | 1.1547005 |
| RIL22 | 5.00 | 3.0000000 | 3.67 | 1.1547005 |
| RIL23 | 1.67 | 1.1547005 | 1.00 | 1.0000000 |
| RIL25 | 3.67 | 1.5275252 | 0.67 | 0.5773503 |
| RIL30 | 1.33 | 1.5275252 | 1.33 | 1.5275252 |
| RIL41 | 1.67 | 1.1547005 | 3.00 | 3.0000000 |
| RIL47 | 4.00 | 0.0000000 | 3.33 | 1.5275252 |
| RIL50 | 5.00 | 2.6457513 | 4.33 | 1.1547005 |
| RIL57 | 2.67 | 0.5773503 | 2.67 | 1.1547005 |
| RIL58 | 5.00 | 3.0000000 | 2.33 | 2.3094011 |
| RIL80 | 8.67 | 0.5773503 | 10.00 | 0.0000000 |
| RIL81 | 2.67 | 1.1547005 | 2.67 | 1.1547005 |
| RIL93 | 6.67 | 0.5773503 | 5.33 | 2.0816660 |

**Table S3.**
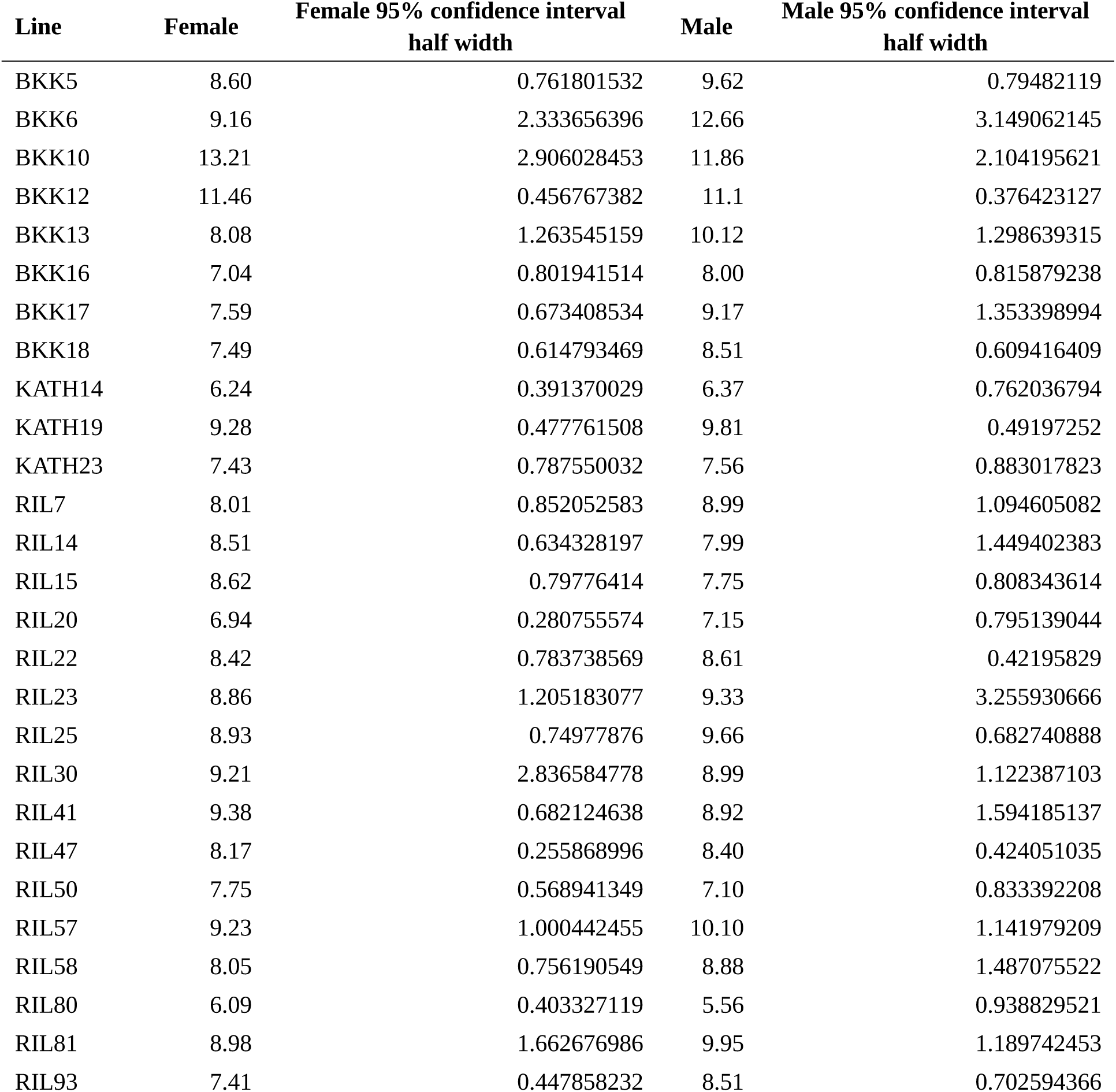
Lethal time to 50% mortality (LTi50, hours) for female and male flies of BKK, KATH, and RIL strains. RIL founder strains are indicated in **bold**.

**Table S4.**
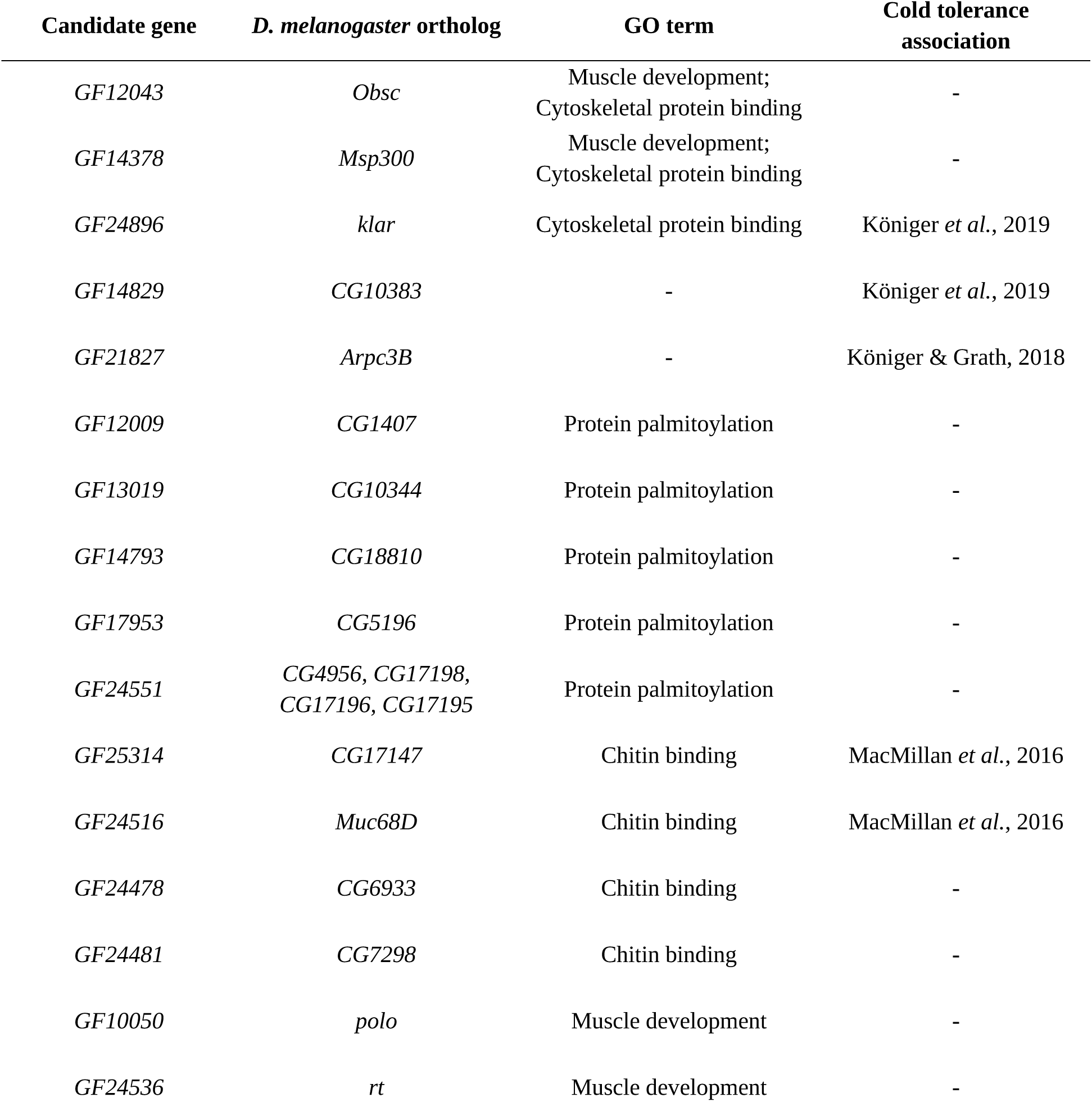
Candidate genes located within the candidate regions identified in the bulk segregant analysis. The table lists *Drosophila ananassae* gene identifiers, annotations, and corresponding *Drosophila melanogaster* orthologs. Gene ontology (GO) enrichment analysis was performed using the *D. melanogaster* orthologs of the candidate genes.

## Supplementary Figures

**Figure S1.**
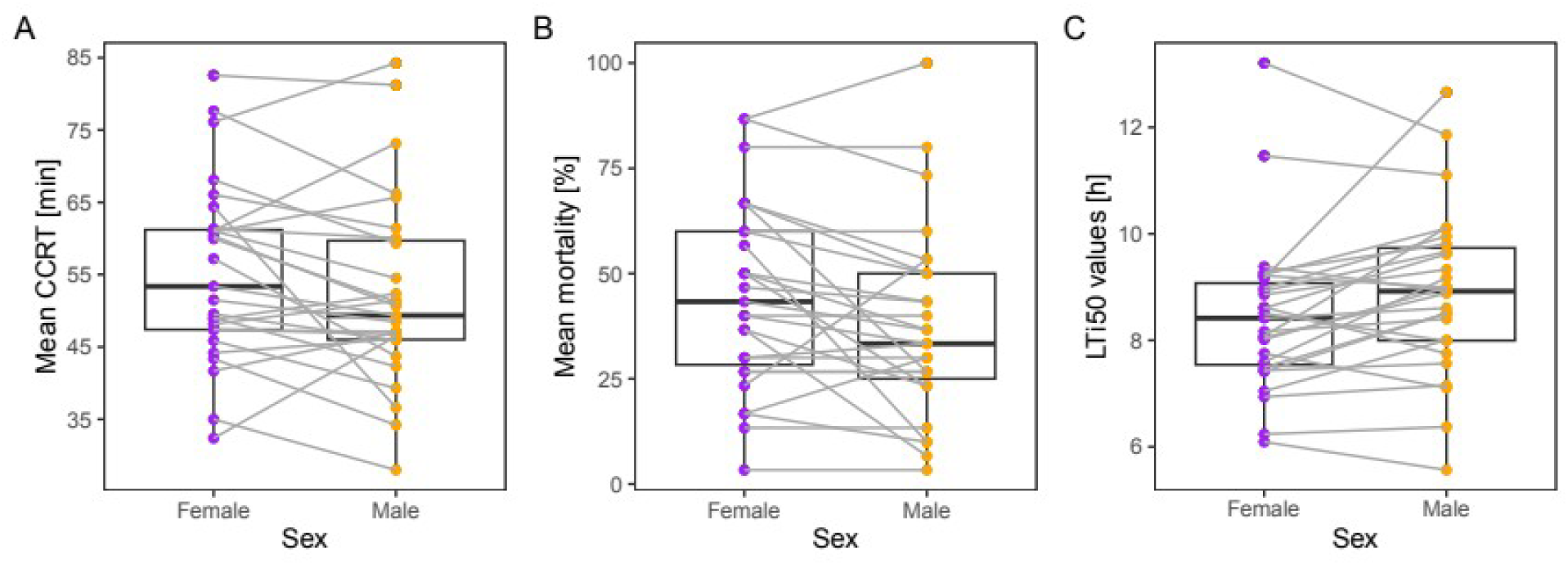
Sex-specific comparison of cold tolerance phenotypes. Boxplots show chill coma recovery time (CCRT, minutes) (A), mortality after an 8 h cold shock (%) (B), and lethal time to 50% mortality (LTi50, hours) (C). Females are shown in purple and males in orange; female and male values from the same strain are connected by grey lines to visualize within-strain differences. Boxes indicate the interquartile range, horizontal lines represent the median, whiskers extend to 1.5× the interquartile range, and outliers are shown as dots. *Alt text: Boxplots comparing female and male CCRT, cold-shock mortality, and LTi50 across strains. Female and male values from the same strain are connected by grey lines, showing the magnitude and direction of sex differences within individual strains*.

**Figure S2.**
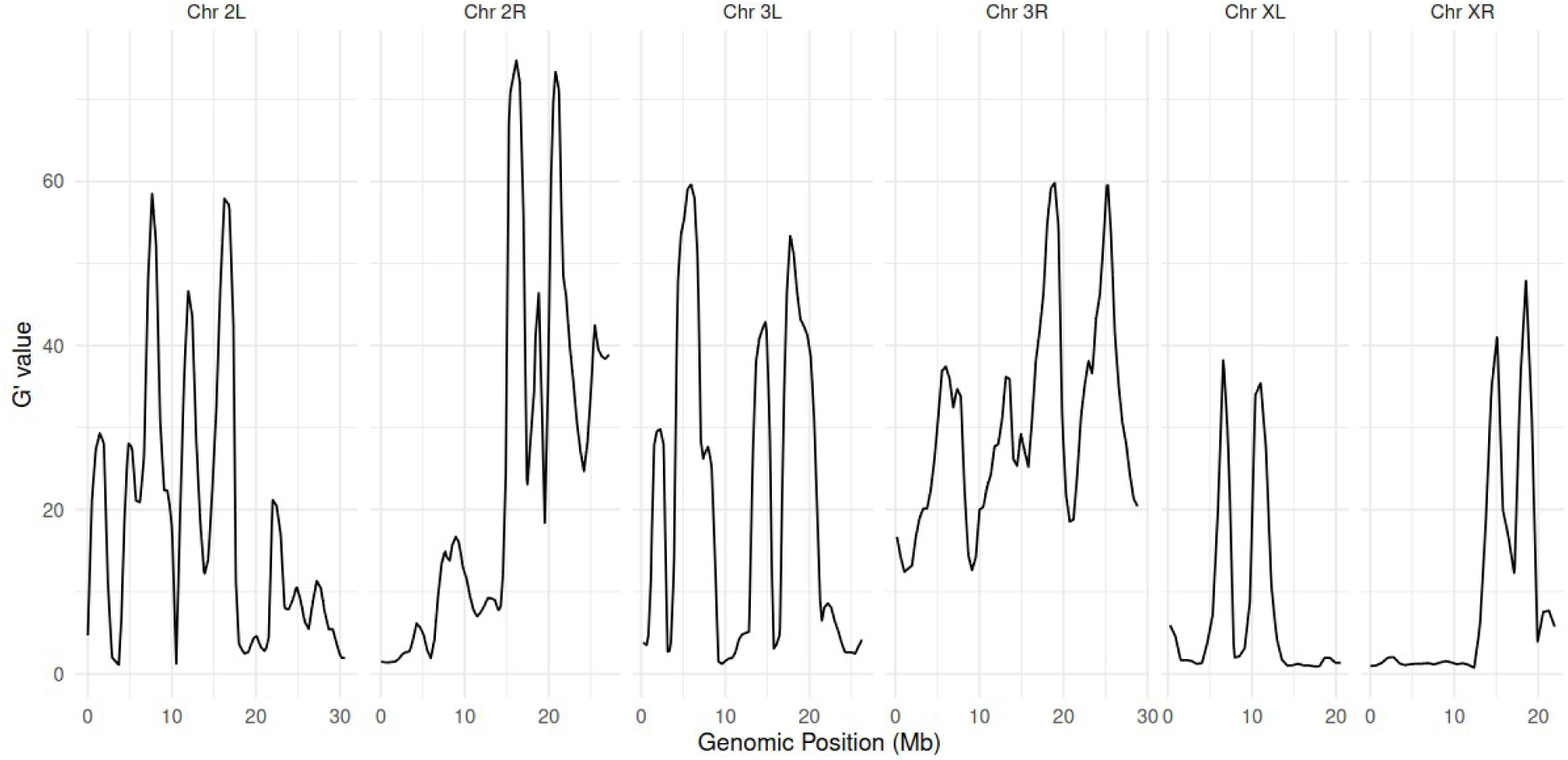
QTL-seq analysis using the G′ statistic. G′ values calculated across the genome within 1 Mb sliding windows. The x-axis shows genomic position and the y-axis shows G′ values. No genomic regions exceeded the significance threshold in this analysis. *Alt text: Genome-wide plot of G′ values across chromosomes showing no regions exceeding significance thresholds, indicating no strong associations detected using this method*.

**Figure S3.**
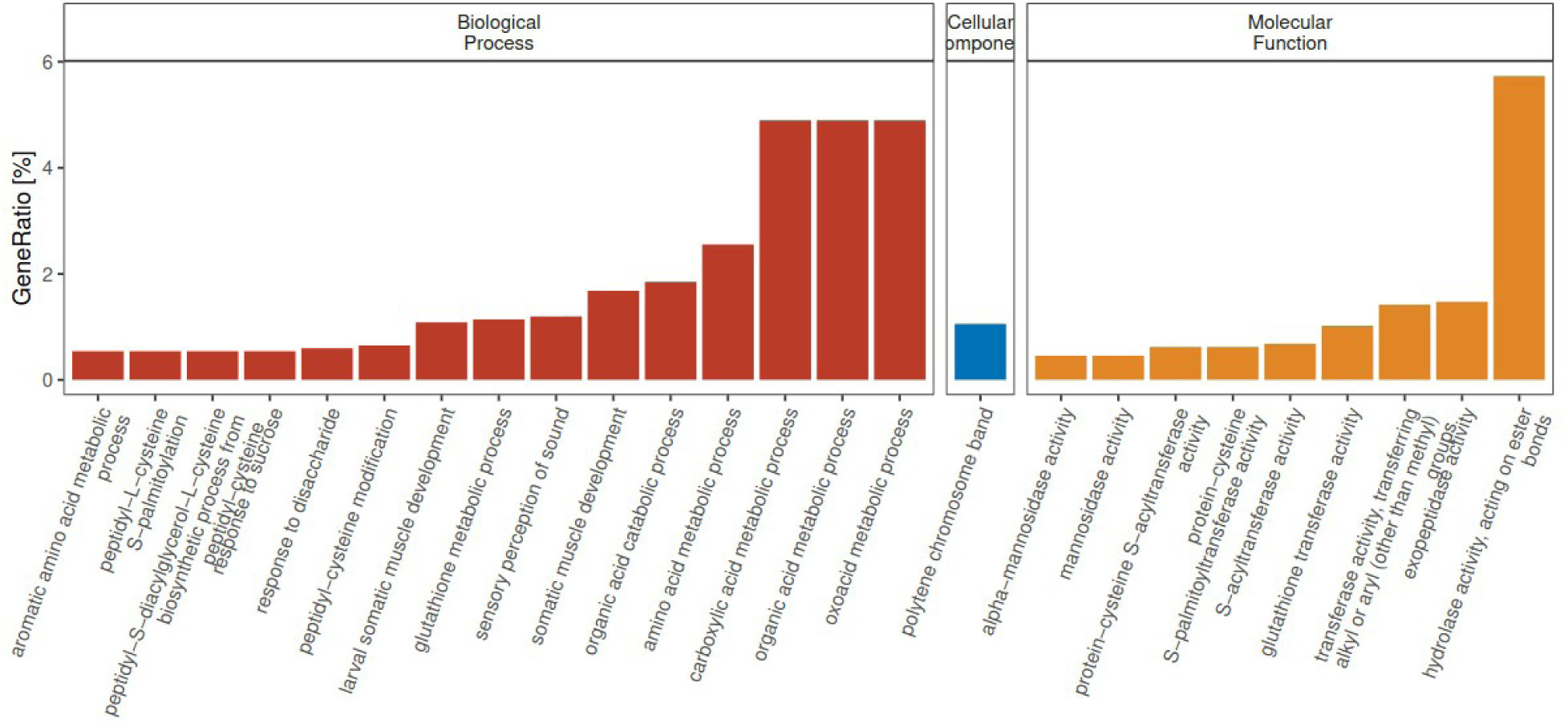
Gene ontology (GO) terms enriched in candidate regions with significant positive Δ(SNP-index) values. The top 25 GO terms ranked by p-value are shown. The y-axis indicates the ratio of genes associated with each GO term. Biological processes are shown in red, cellular components in blue, and molecular functions in yellow. *Alt text: Bar plots of gene ontology enrichment for genes in candidate regions with positive allele frequency differences, highlighting functions related to muscle development, metabolism, and cytoskeletal organization*.

**Figure S4.**
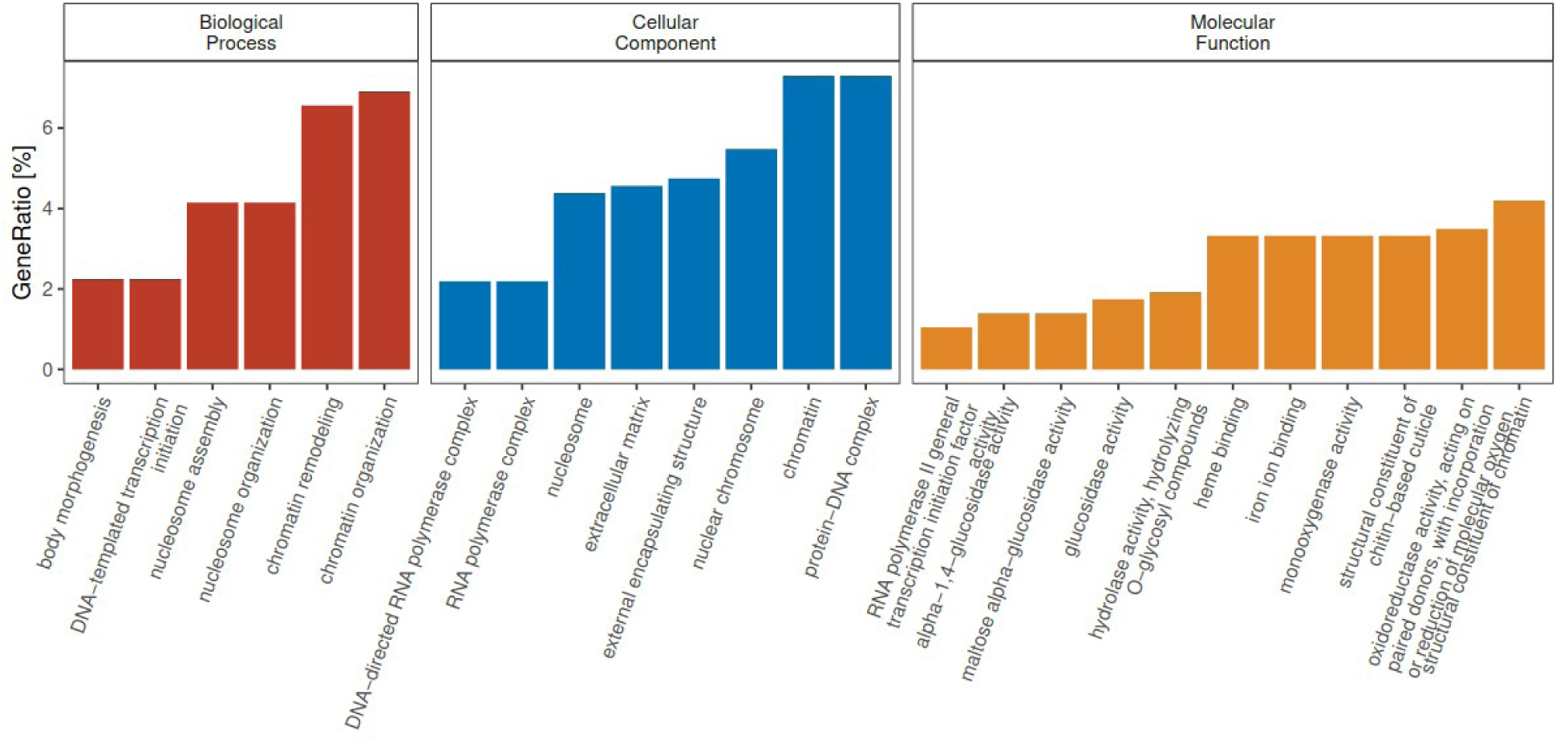
Gene ontology enrichment analysis for genes within the candidate regions with significant negative *Δ(SNP-index) values.* Biological processes are shown in red, cellular components in blue, and molecular functions in yellow. *Alt text: Bar plots of gene ontology enrichment for genes in candidate regions with negative allele frequency differences, showing enrichment of enzymatic activities including glucosidase, monooxygenase, and hydrolase functions*.

## Notes

### Competing Interest Statement

The authors have declared no competing interest.

### Summary of Updates

Revision upon reviewer comments; Figures revised; supplemental files uploaded; text revision

